# Above-Filter Digestion Proteomics reveals drug targets and localizes ligand binding site

**DOI:** 10.1101/2025.03.11.642584

**Authors:** Bohdana Sokolova, Hassan Gharibi, Maryam Jafari, Hezheng Lyu, Silvia Lovera, Massimiliano Gaetani, Amir Ata Saei, Roman A. Zubarev

**Author notes:** Correspondence and requests for materials should be addressed to A.A.S. and R.A.Z.

## Abstract

While a number of efficient chemical proteomics methods are available for determining protein targets of pharmaceutical drugs, each approach exhibits its own “blind spot” and development of complementary techniques is needed. Here, we introduce the Above-Filter Digestion Proteomics (AFDIP) approach based on the monitoring of the rate of trypsin digestion. Digestion rate is decreased at the site of ligand binding to its target protein, while other sites may simultaneously experience an increase the digestion rate. Molecular dynamics (MD) simulations revealed that this increase may be due to allosteric structural changes related to backbone flexibility. We showcase the utility of AFDIP for deconvoluting the targets of versatile drugs and metabolites. Like other techniques, AFDIP allows for a two-dimensional analysis, with the second dimension based on drug concentration. AFDIP exhibits structural resolution, identifying drug target binding sites within ≤10 Å, and for larger proteins often ≤5 Å of the crystallography-determined binding sites. Compared to earlier introduced limited proteolysis approaches, AFDIP provides easier sample preparation, deeper proteome analysis and broader sequence coverage. AFDIP is expected to find its place among the most efficient chemical proteomics methods currently available.

## Introduction

Determination of the drug target is a primary goal in chemical proteomics and is an integral component of every drug development campaign. Several efficient strategies have already been developed to address this problem. Some of these approaches are based on expression proteome profiling, including Functional Identification of Target by Expression Proteomics (FITExP)^1^ and its derivative ProTargetMiner^2^. Thermal Proteome Profiling (TPP) identifies drug targets by assessing the ligand-induced thermal stability changes of cellular proteins^3^, adding the strength of system-wide proteomics to Cellular Thermal Shift Assay (CETSA)^4^. TPP enables sorting out the differences in melting temperatures of proteins due to direct ligand-binding effects in cellular extracts and those resulting from downstream cellular modifications in living cells^3^, and can be used for identification of drug targets and off-targets^5^, tracking target engagement in cells and tissues^6^. The introduction of the Proteome Integral Solubility Alteration (PISA)^7^ assay not only enhanced the throughput compared to TPP by at least tenfold, but also substantially increased the depth of proteome coverage. Due to the capability to analyze both treated and untreated samples in several biological replicates simultaneously, PISA offers enhanced statistical power and requires considerably less sample amount for analysis. Moreover, protein solubility in PISA can be modulated not only with temperature, but also with organic solvents^8^ as well as kosmotropic salts^9^. In the latter incarnations, PISA reached sub-µg level sample consumption per replicate^10^. Another chemical proteomics method, isothermal shift assay (iTSA)^11^, measures protein stability shifts at a single temperature point using fixed incubation time, demonstrating improved efficiency in target identification compared to TPP. This enhanced performance is attributed to iTSA’s simplified experimental design, which requires fewer samples and allows for increased replication, leading to greater statistical power. At the same time, the broad range of protein melting temperatures found in nature^12^ makes selection of a single temperature and incubation time problematic, limiting the applicability of the latter approach.

Every drug target identification technique has its limitations, with certain “blind spots” inherent to each method. For instance, in FITExP some drug targets may not change their abundance strongly enough to be detected with statistical significance, while in PISA the solubility of some proteins may not be altered markedly upon ligand binding. In our experience, each chemical proteomics method provides on average the correct answer in approximately 60-70% of the cases. This necessitates the use of several complementary proteome-wide approaches for achieving high statistical power of target identification, as we have shown in the past^13^. For instance, in order to ensure detection of drug-target interaction in ≥95% cases at a 65% success rate for every approach one needs to combine three orthogonal techniques.

Besides, neither of the aforementioned techniques offers insights into structural detail of drug-target interactions. This objective is achieved, for instance, by the Stability of Proteins from Rates of OXidation (SPROX)^14^ technique that assesses the changes in target protein stability under oxidative conditions. Hydrogen peroxide is used to oxidize the methionine residues in target proteins and the oxidation occupancies are measured in each sample by proteomics, being inversely related to the frequency of drug binding to the target. The disadvantage of SPROX is the relatively low frequency of methionine residues in the proteins. Therefore, while SPROX has been successfully used to identify the targets of small molecules such as cyclosporine A^15^, it is inferior to both FITExP and PISA in terms of the proteome depth of analysis.

Another such complementary method is drug affinity responsive target stability (DARTS)^16^. This approach provides insight into drug-protein interactions based on the fact that drug binding can make proteins more resistant to proteolysis. In DARTS, a protein mixture is incubated with the drug of interest and then subjected to partial proteolysis by thermolysin for a fixed time duration (typically, 10 min). The resulting peptides are analyzed by gel electrophoresis and mass spectrometry, with protected proteins indicating potential drug targets. DARTS has been successfully used to identify targets for various compounds, however, its main limitation is that proteins have susceptibility to proteolysis based on their conformational properties. Therefore, choosing conditions for partial proteolysis which could be optimal at the proteome level for target identification may be problematic. Besides, as thermolysin is an unspecific protease preferentially cleaving at the N-terminal side of hydrophobic or bulky amino side chains such as Leu, Phe, Ile, and Val, the abundance of non-tryptic peptides complicates MS/MS data processing.

A more recent approach based on the same phenomenon is Limited Proteolysis (LiP)^17^. Similar to DARTS, LiP relies on time limited hydrolysis of polypeptide bonds in proteins by unspecific proteases (most commonly, proteinase K), but this is followed by full digestion of the partially hydrolyzed proteins by trypsin and LysC. The mixture of fully tryptic and semi-tryptic peptides is then analyzed by LC-MS/MS. Differences in LiP patterns for treated and untreated samples are then linked to protein structure, enabling the detection of drug-target interaction surface. As most chemical proteomics approaches, LiP has a broad range of applications from drug target identification^18^ to mapping protein-metabolite^18,19^ and protein-protein interactions^20^. However, the statistical analysis in LiP is not trivial. This is mostly because target identification must be performed at the peptide level without the benefit of the common for both expression proteomics and PISA assay assumption that all peptides belonging to the same protein behave the same way. Thus, in order to obtain reliable results, machine learning needs to be employed, as e.g., in LiP-Quant^18^ approach, to detect drug-binding indicative features. These features are then combined into a unified score for identifying the protein targets of small molecules and estimating their binding locations.

The main advantage of LiP and similar proteolysis-based methods to conventional chemical proteomics is that, theoretically, they have the potential of identifying the site of drug-target interaction. The structural resolution of LiP, however, has so far been quite restricted, with binding site identification limited to a protein domain or part of the sequence covering up to half of the protein^18^. This is because, besides direct interactions of the drug with the target protein, LiP is also sensitive to allosteric changes in protein structure, which can happen far away from the binding site. The sequential use of two enzymes as well as limited sequence coverage also reduces the structural resolution.

There are other difficulties in the LiP approach, such as the abundance of semi-tryptic peptides arising due to the nonspecific cleavage by proteinase K. The presence of semi-tryptic peptides greatly expands the search space in peptide sequence identification by MS/MS, and increases either the threshold score for positive identification, false detection rate (FDR) or both^21^. As mentioned, in LiP, only one or a few peptides of the target protein are expected to shift in the presence of the drug. Since the number of peptides identified in a proteomic analysis usually exceeds that of proteins by an order of magnitude exceeds that of proteins, this creates an additional burden arises when performing multiple hypothesis correction to the p-value.

Recently, PELSA^22^ (peptide-centric local stability assay), another proteolysis-based proteomics method for identifying protein targets and binding regions of diverse ligands, has been introduced. Unlike traditional partial-proteolysis approaches, PELSA employs a large amount of trypsin (enzyme-to-substrate ratio of 1:2) to generate small peptides directly from proteins under native conditions. This approach allows for sensitive detection of ligand-induced protein local stability shifts on a proteome-wide scale. However, despite showing a number of spectacular results, PELSA also has some weak spots. For instance, an ultra-short one-minute digestion time may introduce variability between replicate experiments, and shallow digestion results in a lower protein sequence coverage compared to LiP-Quant^18,22^. As identification of the target protein in PELSA is based on a single peptide, low sequence coverage may lead to false negatives, as non-unique peptide may also result in false positives. The high cost of sequence-grade trypsin is another limiting factor.

Despite their limitations, partial proteolysis approaches are promising in chemical proteomics, being complementary to both expression-based techniques (FITExP) and solubility-based strategies (e.g., PISA). To be competitive with these powerful techniques, at least some of the above drawbacks need to be eliminated. Here we present such a technique called Above-Filter Digestion Proteomics (AFDIP). Unlike LiP and similar to PELSA, AFDIP uses exclusively trypsin, but unlike PELSA it achieves full digestion. This approach offers deeper sequence coverage and lower FDR due to the absence of semi-tryptic peptides, while retaining the structural information. Also, the use of trypsin makes AFDIP data processing compatible with other proteomics approaches. While the “contrast” in determining the binding site is lower compared to PELSA, the target protein is identified in AFDIP by several peptides, providing statistically robust protein identification. Here we describe AFDIP in detail and demonstrate that, on the same model systems as LiP, it offers similar if not superior performance without the need to involve machine learning and using conventional statistical approaches with multiple hypotheses correction.

Figure 1a shows the outline of time-AFDIP workflow. The key element is the trypsin digestion of the extracted proteome incubated with either a drug or vehicle that is performed in a vial above a membrane filter with a 3 kDa molecular weight cut-off (MWCO). The extracted cell lysate is divided into two groups: one treated with the drug and the other (control) treated with only the vehicle. Both groups are then incubated with trypsin. With one-hour intervals, the samples are centrifuged on the MWCO filters, after which the filtrate is collected and additional buffer is added to the undigested proteins above the filter. After 8 h of digestion, all collected filtrates representing different time points for control and treated samples are labelled with the isobaric tandem mass tag (TMT). The labeled filtrate fractions are then combined, providing in the case of TMT-16 labeling a single replicate of AFDIP analysis. Multiplexing into one TMT set the samples obtained with and without the drug reduces the missing value problem and increases the analysis precision. Such a procedure is performed for at least three replicates.

**Figure 1.**
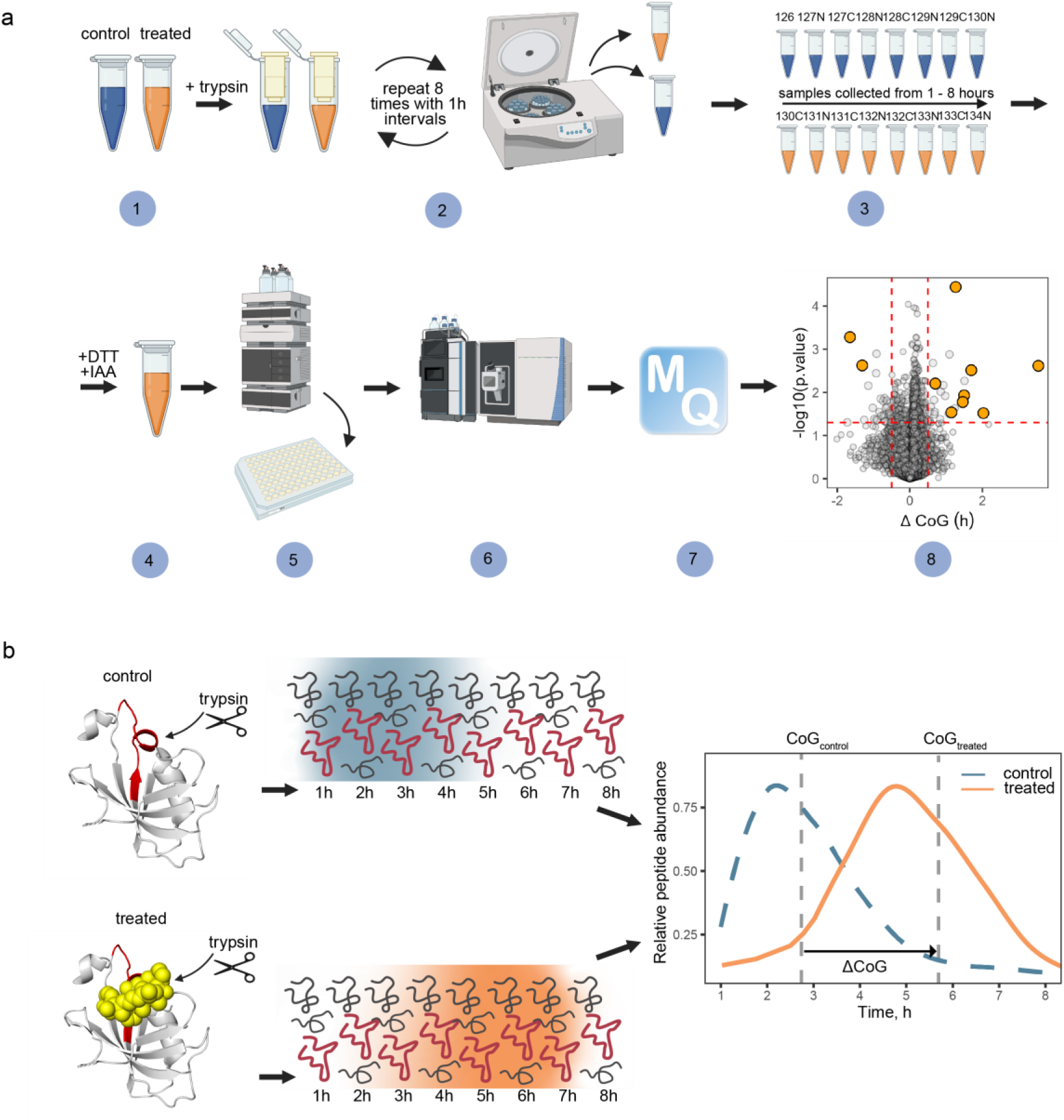
Schematics and workflow of AFDIP. (a) Time-AFDIP experimental workflow. After drug treatment, trypsin is added to all samples (1). After every hour of digestion, the samples are loaded on 3 kDa MWCO filters and centrifuged. The filtrate is collected, and buffer is added to the undigested proteins; the procedure is repeated for 8 h (2). Collected filtrate from each timepoint for control and treated samples is labelled with a TMT (3). All timepoint samples for one replicate are combined, DTT and IAA are added to reduce and alkylate cysteine residues (4). Peptides are separated by hydrophobicity into fractions by HPLC (5). Samples are analyzed with LC-MS/MS (6), then a database search and data analysis are performed (7), resulting in a volcano plot (8). (b) Generation of the peptide yield curves and quantification of the Center of gravity shift (ΔCoG).

For deeper proteomics analysis, the multiplexed TMT-labeled sample can be fractionated by reversed phase liquid chromatography into 8-32 fractions. After LC-MS/MS analysis of the fractions, each peptide is found to produce a bell-shape elution curve shown in Figure 1B.

As the time-AFDIP readout, the center of gravity for each peptide is calculated as:

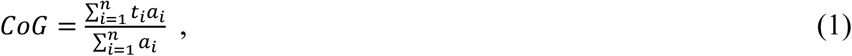

where CoG stands for the center of gravity, *t* indicates a timepoint in hours (1-8), and *a* is the normalized relative abundance of a peptide of interest at a given timepoint. The shift ΔCoG is then determined as the difference between CoG when the drug is present compared to vehicle-only sample. The p-value for the shift is calculated for each peptide based on the replicate results. The result is then presented in a volcano plot, with statistically significant outliers investigated as potential drug targets (Figure 1a). We also test the power of AFDIP in structural analysis of drug-target interactions and demonstrate that it is at least equivalent to that in LiP.

We first chose methotrexate (MTX) in the lysate of HeLa cells, a commonly used model system in our lab as a proof of principle for new chemical proteomics methods^1,7^, as MTX binds strongly to the protein DHFR in both living cells as well as in lysates. After achieving a proof of principle, we investigated the dose-response effect (conc-AFDIP) as second dimension for drug target elucidation at the proteome-wide level.

Another model system employed in HeLa cell lysate was the mTOR inhibitor rapamycin, also studied by LiP^18^. To validate the time-AFDIP rapamycin findings, we additionally analyzed tacrolimus (also known as Fujimycin or FK506), which according to literature has the same main target (FK506-binding proteins) and binding site as rapamycin^23^. Besides the drug targets, we investigated the structural resolution of time-AFDIP and compared it with that of LiP. To put the results of time-AFDIP in a structural context, we compared them with the results of molecular dynamic simulations. Finally, we tested time-AFDIP on promiscuous kinase inhibitor staurosporine^24^ to evaluate its performance compared to other chemoproteomics methods.

## METHODS

### Cell work and proteome extraction

HeLa cells (CCL-2) obtained from ATCC were grown at 37 °C in 5% CO_2_ in DMEM media supplemented with 2 mM L-Glutamine, 100 U/mL Pen/Strep, and 10% FBS.

For proteome extraction, HeLa cell pellets were resuspended in 20 mM EPPS buffer at pH 8. Cells were lysed by 5 freeze-thaw cycles in liquid nitrogen, and the lysate was cleared by centrifugation at 14,000 g for 10 min. Protein concentration was then measured using a BCA Protein Assay kit. 0.5 mL of lysate at the final concentration of 1 mg/mL of protein was aliquoted for further analysis.

### Sample preparation

Lysate samples were either incubated with vehicle (DMSO) or drugs at 10 µM concentration for 30 min at 25°C. For the digestion procedure, trypsin was added at a 1:100 w/w ratio. After every hour of digestion at 25°C over the 8 h timeframe, samples were loaded on a 3 kDa MWCO filter and centrifuged at 14,000 g for 10 min. The filtrate was collected, and buffer was added to the undigested protein above the filter. TMT16 reagents were added 4x by weight to each sample, followed by incubation for 2 h at room temperature (RT). The reaction was quenched by addition of 0.5% hydroxylamine. Samples within each TMT set were combined and reduced with DTT (final concentration10 mM) for 1 h at RT. Afterwards, iodoacetamide (IAA) was added to a final concentration of 50 mM followed by 1 h incubation in the dark. Samples were then acidified by trifluoroacetic acid (TFA), cleaned using Sep-Pak and dried using a DNA 120 SpeedVac™ concentrator (Thermo). The pooled samples were resuspended in 20 mM ammonium hydroxide and separated into 96 fractions on an XBrigde BEH C18 2.1 × 150 mm column (Waters; Cat#186003023), using a Dionex Ultimate 3000 2DLC system (Thermo Scientific) over a 48 min gradient of 1–63% B (B = 20 mM ammonium hydroxide in ACN) in three steps (1–23.5% B in 42 min, 23.5–54% B in 4 min and then 54–63%B in 2 min) at 200 µL min^−1^ flow. Fractions were then concatenated into 24 samples in sequential order (e.g. 1, 25, 49, 73). After resuspension in 0.1% FA (Fisher Scientific) to a concentration of 1 µg/ µL, each fraction was analyzed by LC-MS/MS.

For the dose response experiment, lysate samples were aliquoted into 18 tubes at a final protein concentration of 1 mg/mL. Samples were then treated with either vehicle (DMSO) or methotrexate at concentrations ranging from 0 to 1000 nM and incubated for 30 min at 25°C. Trypsin was added to each sample at a 1:100 ratio, followed by incubation at 25°C for 2 h after which the digestion was inhibited by adding aprotinin. TMT reagents were added to each sample at a 4x weight ratio. After the 2 h long incubation at 25°C, the reaction was quenched by adding 50% hydroxylamine to a final concentration of 0.5%.

Individually labeled TMT samples in each replicate were pooled and loaded onto the filter units and centrifuged at 14,000 g for 10 min, with the filtrate (containing peptides in EPPS) retained for further processing. DTT was added to a final concentration of 10 mM, followed by incubation at 37°C for 30 min. IAA was added to a final concentration of 50 mM and incubated for 30 min at RT in the dark. Samples were acidified with TFA to pH 2-3 and cleaned using SepPak. After drying, samples were fractionated like above into 24 fractions.

### LC-MS/MS analysis

Samples (ca. 1 µg) were loaded onto a 50 cm column (EASY-Spray, 75 µm internal diameter (ID), PepMap C18, 2 µm beads, 100 Å pore size) connected to an Easy-nLC 1000 pump (Thermo Fisher Scientific) and eluted with a 150-min gradient reaching from 3% to 95% of buffer B (98% ACN, 0.1% FA, 2% H_2_O) in buffer A (0.1% FA, 2%ACN, 98% H_2_O) at a flow rate of 300 nL/min. Mass spectra were acquired with an Orbitrap Fusion Lumos Tribrid mass spectrometer (Thermo Fisher Scientific) in the data-dependent acquisition (DDA) mode at a nominal resolution of 120,000 for MS and 45,000 for MS/MS, in the m/z range from 375 to 1500. Peptide fragmentation was performed via higher-energy collision dissociation (HCD) with normalized energy set at 33%.

### Data processing

The raw data from mass spectrometry was analyzed by MaxQuant, version 1.5.3.8^25^. The Andromeda search engine^26^ searched MS/MS data against the International Protein Index database (human version 2014_02, 89,054 entries). Cysteine carbamidomethylation was used as a fixed modification, whereas N, Q-deamidation was selected as a variable modification. Trypsin/P was selected as enzyme specificity. No more than two missed cleavages were allowed. For all other parameters, the default settings were used.

### Statistical analysis

Data was processed by Excel, R and Python. Data for each peptide was analysed using a custom R script that processed intensity values across an 8-hour time course with for both treated and control conditions (n=3 replicates). Intensity values were normalized to the maximum intensity within each condition, and CoG shifts were quantified by calculating weighted means (centroids) for each profile, with statistical significance assessed by two-sided unpaired Student’s t-tests of centroid positions between conditions (p < 0.05), while temporal patterns were modelled using fourth-order polynomial regression.

### Target protein identification

To identify the drug targets of the small molecules, the p-values for peptides originating from the same protein were combined using a Fisher’s combined probability test^27^. This method transforms the sum of the natural logarithms of p-values into a chi-squared statistic (*X_2k_*^2^ = −2 ∑_i=1_^*k*^ ln p_*i*_) that follows a chi-squared distribution with 2k degrees of freedom, where k is the number of p-values being combined. The probability derived from this distribution represents the combined significance at the protein level.

### Binding site identification

The ability of AFDIP to identify binding sites was assessed with PyMOL^28^ software. For each system, peptides of the target protein were sorted by their p-values, and top three peptides were selected for the binding site elucidation. Then, the center of mass (CoM) of all atoms belonging to these peptides was calculated. As a reference, CoM of all residues within 5 Å distance of a ligand was used.

### Molecular dynamic simulations

Molecular dynamics simulations were performed using OpenMM^29^ using the Amber ff14SB force field^30^. Force field parameters for the FK506 ligand were based on the general Amber force field (Gaff)^31^. Long-range electrostatics were treated with particle mesh Ewald summation (PME)^32^. Hydrogen atoms were constrained using the H++ algorithm^33^. Starting structures for the simulations of apo and FK506-bound FKBP52 were taken from PDB entries 1Q1C^34^ and 4LAX^35^, respectively. The simulations were performed for 1 µs in triplicates for both apo and holo structures. Trajectories were then analysed with MDAnalysis^36^.

### Data availability

The LC-MS/MS raw data files and extracted peptides and protein abundances are deposited in the PRIDE repository of the ProteomeXchange Consortium^37^ under the dataset identifier PXD061498.

## RESULTS

### MTX – time-AFDIP

The time-AFDIP volcano plot for the MTX treatment and p-values is shown in Figure 2a. In total, 92,326 peptides belonging to 8001 proteins were quantified in AFDIP analysis. Of these, 18 peptides belonged to DHFR, and covered 85.6 % of the protein sequence. Five peptides showed significant ΔCoG shifts; their AFDIP plots are shown in Figure 2 c. None of the DHFR peptides demonstrated a significant negative shift. When the peptides were sorted by the minimum sum of their ranks of the absolute values of ΔCoG and log10(p), all five of the above mentioned DHFR peptides were among the top 11 peptides, with two peptides receiving the top rankings. For the statistical model of identifying the drug target, the p-values for individual peptides were combined using a Fisher’s combined probability test. When all proteins were ranked by their p-value, DHFR appeared at the top of the list, with p = 7 × 10⁻⁷ (Figure 2b) which even after the stringent Bonferroni correction for multiple hypotheses yielded a significant hit with p* < 6× 10⁻^3^. This result provided a proof of principle demonstrating that AFDIP is capable of identifying the drug target.

**Figure 2.**
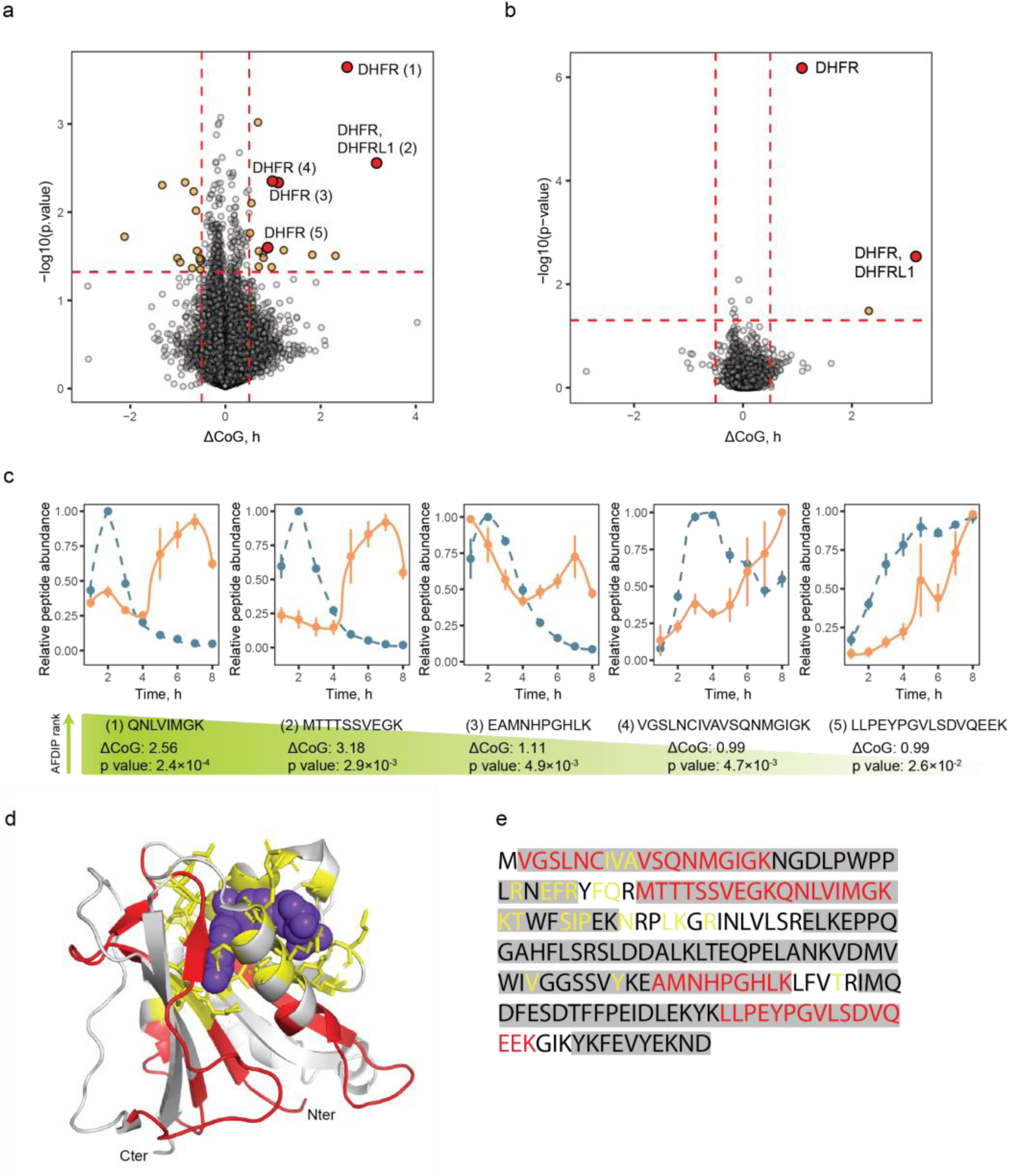
Time-AFDIP of MTX in HeLa lysate. (a) Volcano plot showing in orange the peptides with a significant center of gravity shift (cut-off values are set at 0.5 and-0.5 for ΔCoG and 0.05 for p-value), and peptides from the known MTX target DHFR in red. (b) Volcano plot showing in orange the proteins with a significant center of gravity shift (cut-off values are set at 0.5 and-0.5 for ΔCoG and 0.05 for p-value), and the known MTX target DHFR in red. (c) Center of gravity plots for the shifting peptides of DHFR. Relative abundance of the peptide of interest is plotted for control and treated samples in blue and in orange respectively. (d) Binding site identification with AFDIP. MTX molecule is shown in purple blue, shifting peptides in AFDIP are colored in red, amino acid residues withing 5 Å of the binding site are shown in licorice and colored in yellow. (e) Sequence coverage of DHFR in AFDIP. Peptides detected unshifted are highlighted in gray, shifting peptides are colored in red, and amino acid residues withing 5 Å of the binding site are colored yellow.

We also attempted to obtain insight into the structural details on binding. But as DHFR is a small protein (MW 22 kDa^38^), a significant fraction of its sequence is found close to the canonical binding site with MTX^39^ (Figure 2d). Yet, there was some qualitative correlation between ΔCoG of the peptides and their proximity to the binding site (Figure 2d). For instance, the central part of the protein, not involved in binding, does not contain significantly shifting peptides. To evaluate quantitatively the capacity of AFDIP for binding site elucidation, we calculated the distance ΔCoM between the center of mass (CoM) of all atoms of the binding site (residues within 5 Å of the ligand) and CoM of the atoms belonging to the 3 peptides with the most significant shifts. The obtained value of ΔCoM=8.3 Å can be compared to the longitudinal molecular length of MTX, which is 21.2 Å^40^. In contrast, ΔCoM for the three peptides showing least significant shifts was 9.2 Å. This result indicates that there is a potential for extracting structural information on drug binding site from the AFDIP data, albeit with a lower resolution than in hydrogen-deuterium exchange mass spectrometry^41^.

### MTX - Dose-response effect (conc-AFDIP)

To investigate the potential of conc-AFDIP as an additional dimension for drug target elucidation, protein lysate samples were exposed to an MTX concentration series of from 0.1 nM to 1000 nM, as well as 0 nM, followed by 2 h digestion with trypsin. Upon compound binding, the yields of proteolytic fragments near the place of drug binding are expected to change, with true target peptides exhibiting a sigmoid dependence on drug concentration. In total, 81,313 peptides from 8306 protein groups were quantified in this experiment. Foldchange was calculated by dividing the average relative peptide abundance of 2 lowest treatment concentrations by 2 highest treatment concentrations (Figure 3a). The top 30,000 peptides with highest peptide abundances were ranked by their p-values and absolute values of log2(foldchange), and in the list of top 10 peptides resulting from such ranking 4 belonged to dihydrofolate reductase. Peptides belonging to DHFR demonstrate lower relative peptide abundances with the increase of drug concentration (Figure 3b), which could be explained by steric hindrance for trypsin digestion caused by drug binding. Our statistical analysis performed similar to time-AFDIP produced the p-value from Fisher’s combined probability test for DHFR of 3 × 10^⁻13^, or, after Bonferroni correction for multiple hypotheses, <3 × 10^⁻9^. While in terms of the p-value conc-AFDIP was superior compared to time-AFDIP, this may be the result of our fortunate choice of the digestion time. To avoid the uncertainty associated with making this choice, the rest of the experiments were performed using time-AFDIP.

**Figure 3.**
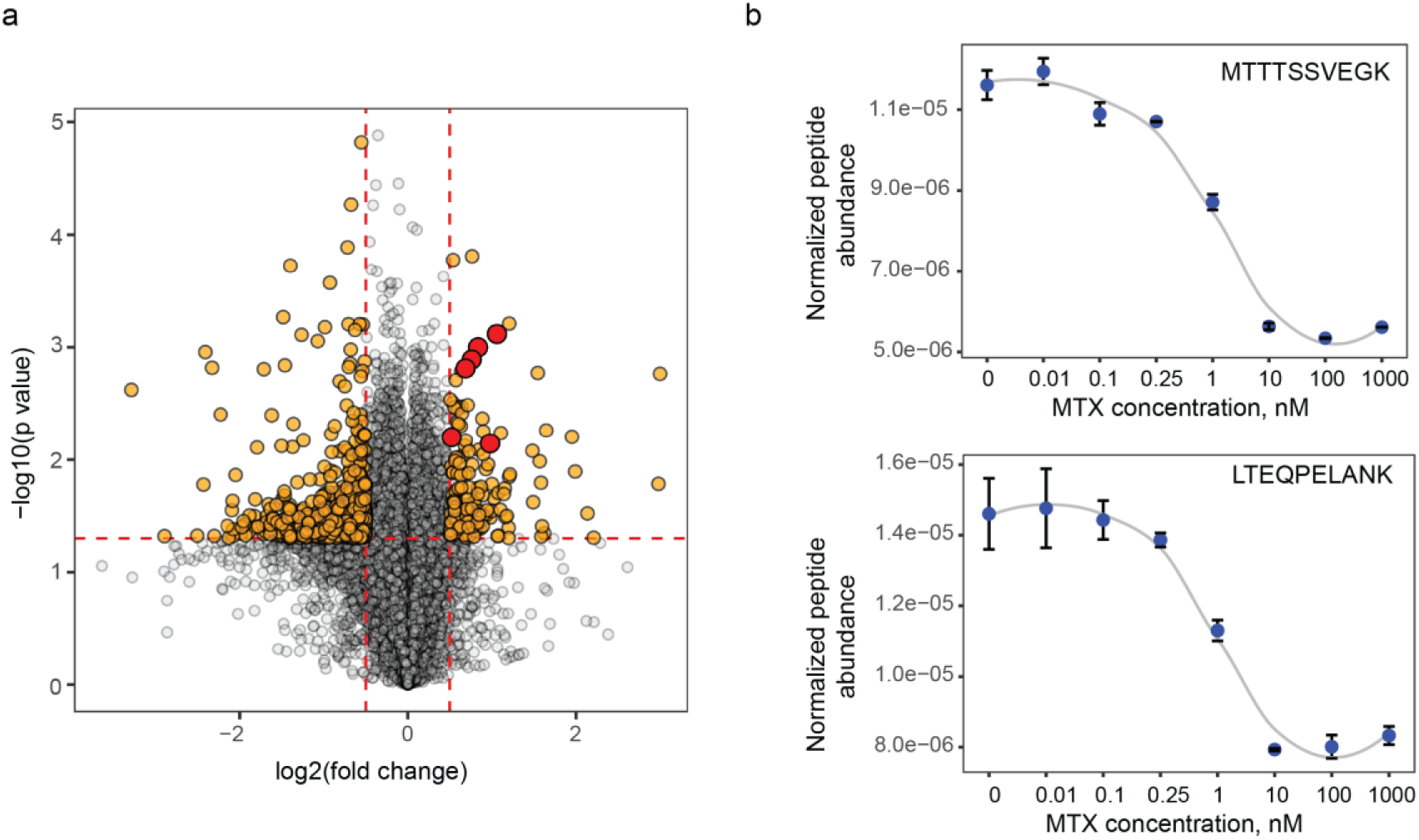
Conc-AFDIP of MTX in HeLa lysate. (a) Volcano plot showing in orange the peptides with a significant foldchange upon drug binding (same cut-offs as in Figure 2), and peptides from DHFR in red. (b) Dose–response curves showing normalized abundances of DHFR peptides as a function of MTX concentration.

### Rapamycin

We then applied AFDIP on rapamycin, an immunosuppressant inhibiting mTOR via complex formation with FK506-binding proteins (FKBPs)^42^. AFDIP analysis quantified a total of 92,286 peptides belonging to 7845 proteins. Among peptides with significant p <0.05 and ΔCoG>±0.5 h, 15 molecules belonged to mTOR and FK506-binding proteins, namely FKBP2, FKBP3, FKBP4, and FKBP5 (see volcano plot on Figure 4a). When the peptides were sorted by the minimum sum of their ranks by absolute values of ΔCoG and log10(p), 8 among the top 10 peptides belonged to FKBPs. FKBP3 peptides were the most numerous (eight) among the 55 significantly shifting peptides. Statistical analysis performed to identify drug target showed that the p-value from Fisher’s combined probability test for FKBP3 was 2.2 × 10^⁻7^, still significant (p < 1.7 × 10^⁻3^) after Bonferroni correction.

**Figure 4.**
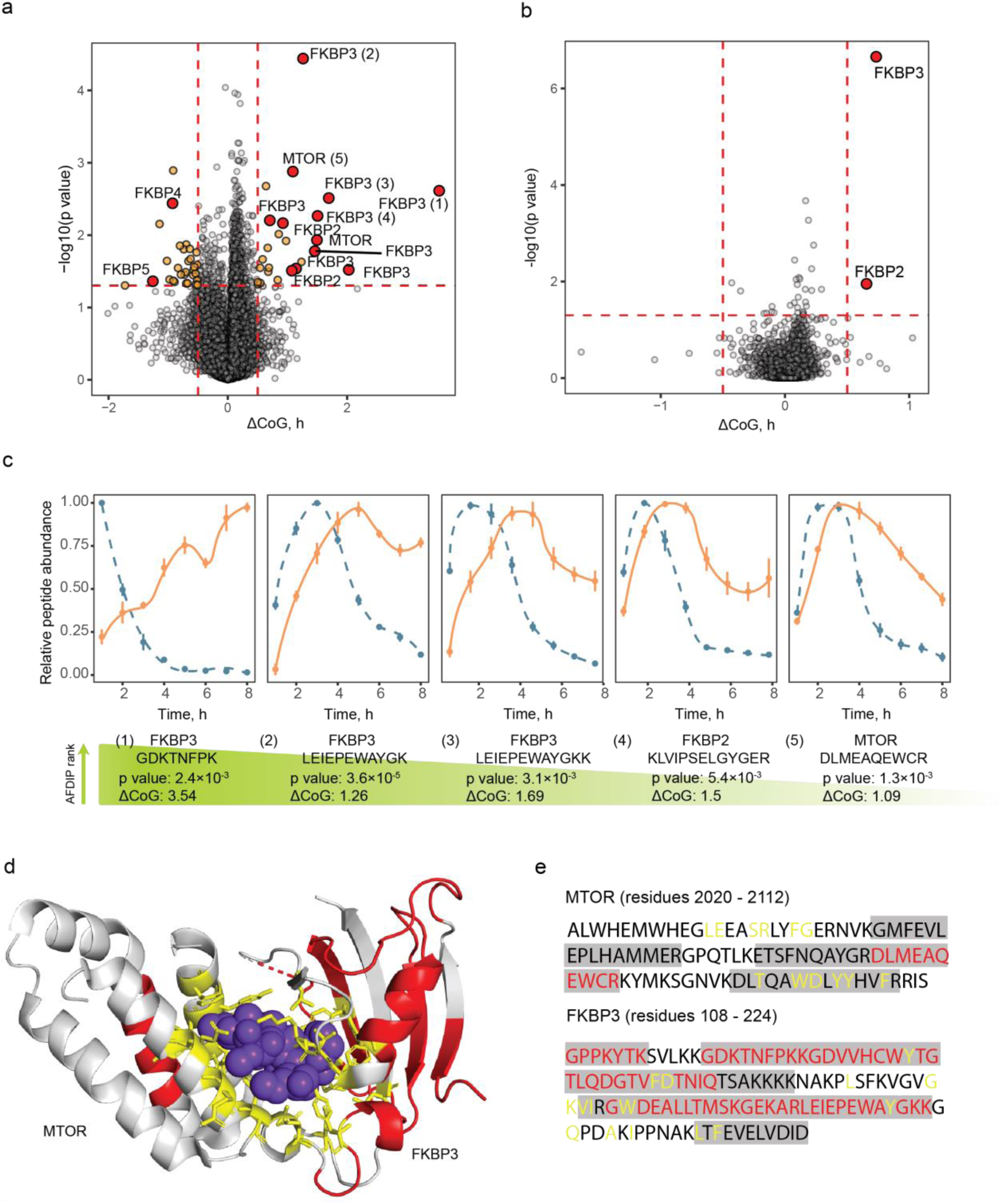
Time-AFDIP of Rapamycin. (a) Volcano plot showing in orange the peptides with a significant center of gravity shift (same cut-off values as in Figures 2 and 3), and peptides from rapamycin’s known targets MTOR, FKBP2, FKBP3, FKBP4, and FKBP5 in red. (b) Volcano plot showing the proteins with a significant center of gravity shift. (c) Yield curves for the shifting peptides of the targets. Control and treated samples are shown in blue and orange, respectively. (d) Binding site identification with AFDIP on rapamycin complex with MTOR and FKBP3 (PDB: 5GPG). Rapamycin molecule is shown in purple blue, shifting peptides in AFDIP colored in red, amino acid residues withing 5 Å of the binding site are shown in licorice and colored in yellow. (e) Sequence coverage of MTOR and FKBP3. Unshifting peptides are highlighted in gray, shifting peptides are colored in red, and amino acid residues withing 5 Å of the binding site are colored in yellow.

Rapamycin together with FKBP3 forms a complex with mTOR ^24^, and consistent with that two mTOR peptides were found among the most significantly shifting peptides (Figure 4). We investigated the capacity of time-AFDIP to identify binding sites on FKBP3-rapamycin-mTOR complex. The distance between CoM of the binding site and top 3 shifting peptides of the complex (of which 2 belonged to FKBP3 and 1 to mTOR) was 3.5 Å. Note that the complex size is quite large (characteristic dimension ≈200 Å^43^) as MW of FKBP3^24^ and MTOR^44^ are 25 kDa and 289 kDa, respectively. Interestingly, the first structure of an mTOR complex was determined by cryogenic electron microscopy (cryo-EM) with a resolution of 4.4 Å^43^. The application of AFDIP to large protein complexes, even in the context of full cell lysate, may therefore potentially compete with this technique in binding site determination.

### FK506

FK506, also known as tacrolimus, is a macrolide immunosuppressant drug primarily used to prevent organ rejection in transplant patients^45^. The FK506 experiment was performed to further explore AFDIP findings with rapamycin, as both drugs bind to FKBPs at the same site yet induce different mechanisms of action in cells. While rapamycin has mTOR as its end target, FK506 forms a complex with FKBPs eventually inhibiting calcineurin/calmodulin pathway^46^.

Volcano plot for AFDIP experiment with FK506 is shown on Figure 5a. Altogether, 109,615 peptides belonging to 8540 proteins were quantified with time-AFDIP. Among the peptides with significant p<0.05 and ΔCoG>±0.5 h, eight molecules belonged to FK506-binding proteins, FKBP2, FKBP4, and FKBP5. When the peptides were sorted by the sum of their ranks of the absolute values of ΔCoG and log10(p), 3 peptides among the top 10 belonged to FKBP2. The p-value from Fisher’s combined probability test for FKBP2 was 1.1 × 10^⁻6^, or p* = 0.01 after Bonferroni correction.

**Figure 5.**
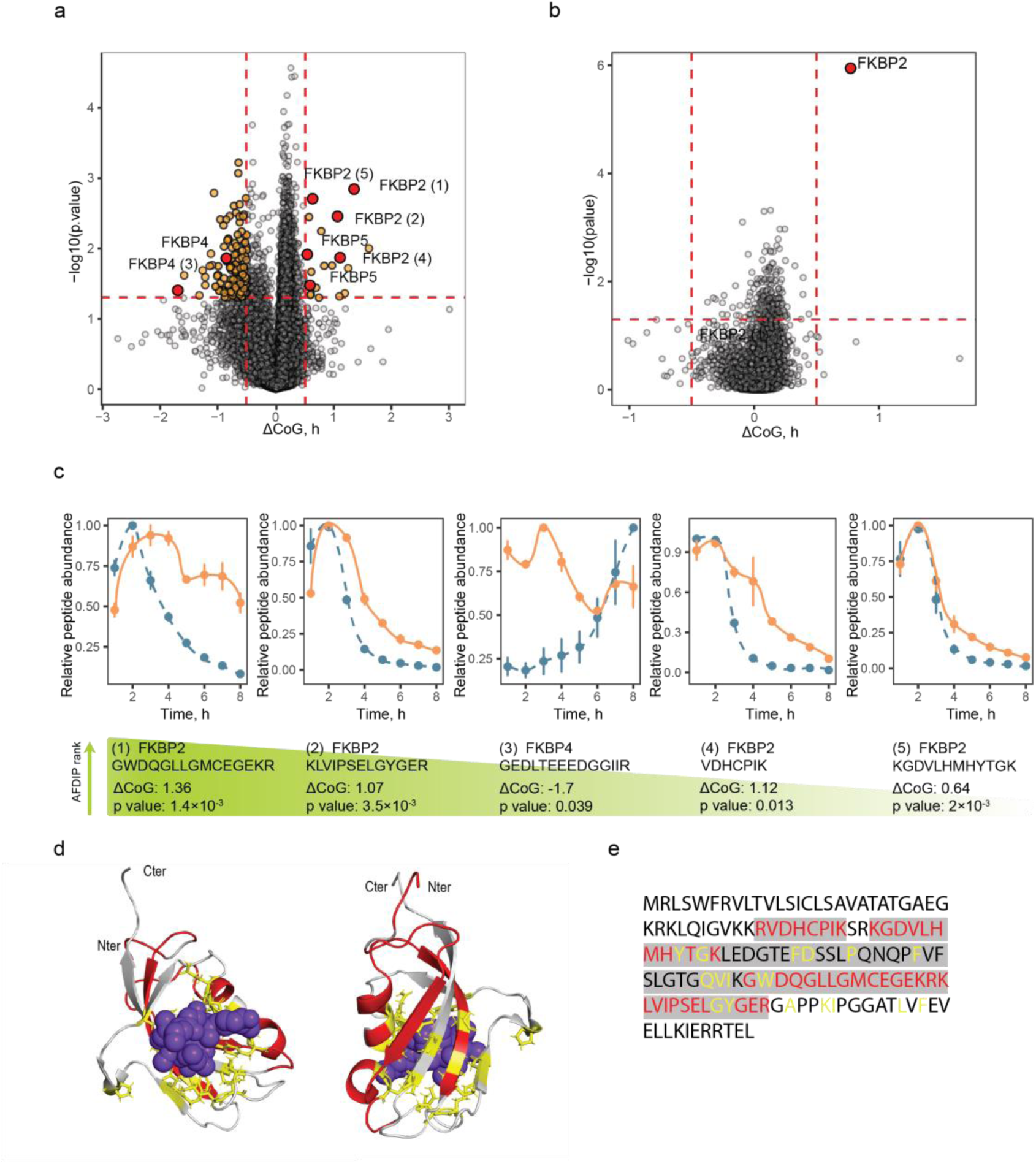
AFDIP of FK506. (a) Volcano plot showing in orange the peptides with a significant center of gravity shift (cut-off values are the same as in Figures 2-4), and peptides from FK506’s known targets FKBP2, FKBP4, and FKBP5 in red. (b) Volcano plot showing the proteins with a significant center of gravity shift. (c) Yield curves for DHFR shifting peptides. Control and treated samples are shown in blue and in orange, respectively. (d) Binding site identification with AFDIP on FK506 complex with FKBP2 (PDB: 4NNR). FK506 molecule is shown in purple blue, shifting peptides in AFDIP colored in red, amino acid residues withing 5 Å of the binding site are shown in licorice and colored in yellow. (e) Sequence coverage of FKBP2. Unshifting peptides detected with MS are highlighted in gray, shifting peptides are colored in red, and amino acid residues withing 5 Å of the binding site are colored in yellow.

Similar to the MTX case, FKBP2 is a relatively small protein (MW 13kDa) with the binding pocket constituting a significant part of the protein sequence. The difference between the average position of the three peptides with the lowest p-value and the binding site from crystallography^47^ was 7.9 Å.

AFDIP was capable of distinguishing between the mechanisms of action of rapamycin and FK506. Both drugs primarily target proteins from the FKBP family yet have varying affinities for FKBPs and different action modes. For instance, FKBP3 shows a significantly higher affinity for rapamycin (Ki 0.9 nM) compared to FK506 (Ki 200 nM)^48,49^. This correlates with AFDIP findings: FKBP3 was not identified as a target of FK506, while being a top target in the rapamycin experiment. Furthermore, mTOR, the downstream target of rapamycin complexes with FKBPs^24^, was exclusively identified as a target in the rapamycin experiment, consistent with its known mechanism of action. Two FKBP2 peptides showing a significant CoG shift in rapamycin experiment were also found among three FKBP2 peptides with the lowest p-value in the FK506 experiment used for the binding site mapping. This demonstrates AFDIP’s capability to identify the common binding site in FKBPs for rapamycin and FK506^49^.

The fact that AFDIP successfully captured the nuanced differences between rapamycin and FK506, while also identifying their common binding site on FKBP2 demonstrates AFDIP’s potential as a powerful tool for discerning subtle variations in drug action mechanisms for compounds sharing similar target proteins.

### MD simulations of FKBP4 with FK506

In the volcano plot of Figure 5a, two FKBP4 peptides shift toward shorter digestion times as opposed to six peptides from other FKBP-family proteins predictively shifting to longer CoGs. In order to investigate this puzzling behavior, we performed MD simulations of the FK1 and FK2 domains of FKBP4 with (holo) and without (apo) the FK506 ligand. Figure 6a shows the root mean square fluctuation (RMSF) of different parts of the protein for both forms. RMSF reflects individual residue’s flexibility, or how much a particular residue moves (fluctuates) in the course of a simulation. Upon normalization, the difference in RMSF values (Figure 6b) highlights the residues with increased (positive ΔRMSF; negative ΔCoG) or decreased (negative ΔRMSF; positive ΔCoG) flexibility upon ligand binding. The positive relative ΔRMSF peak (red band in Figure 6b) corresponds to the flexible linker between FK1 and FK2 domains (Figure 6c) showing the most significant negative ΔCoG. Another strong positive ΔRMSF peak (orange band in Figure 6b) correlates with flexible unstructured region in Figure 6c, which also has a negative ΔCoG in AFDIP. We concluded that the center of gravity shift of a peptide in time-AFDIP may correlate with the flexibility of protein structure that modulates the accessibility to trypsin and its rate of cleavage.

**Figure 6.**
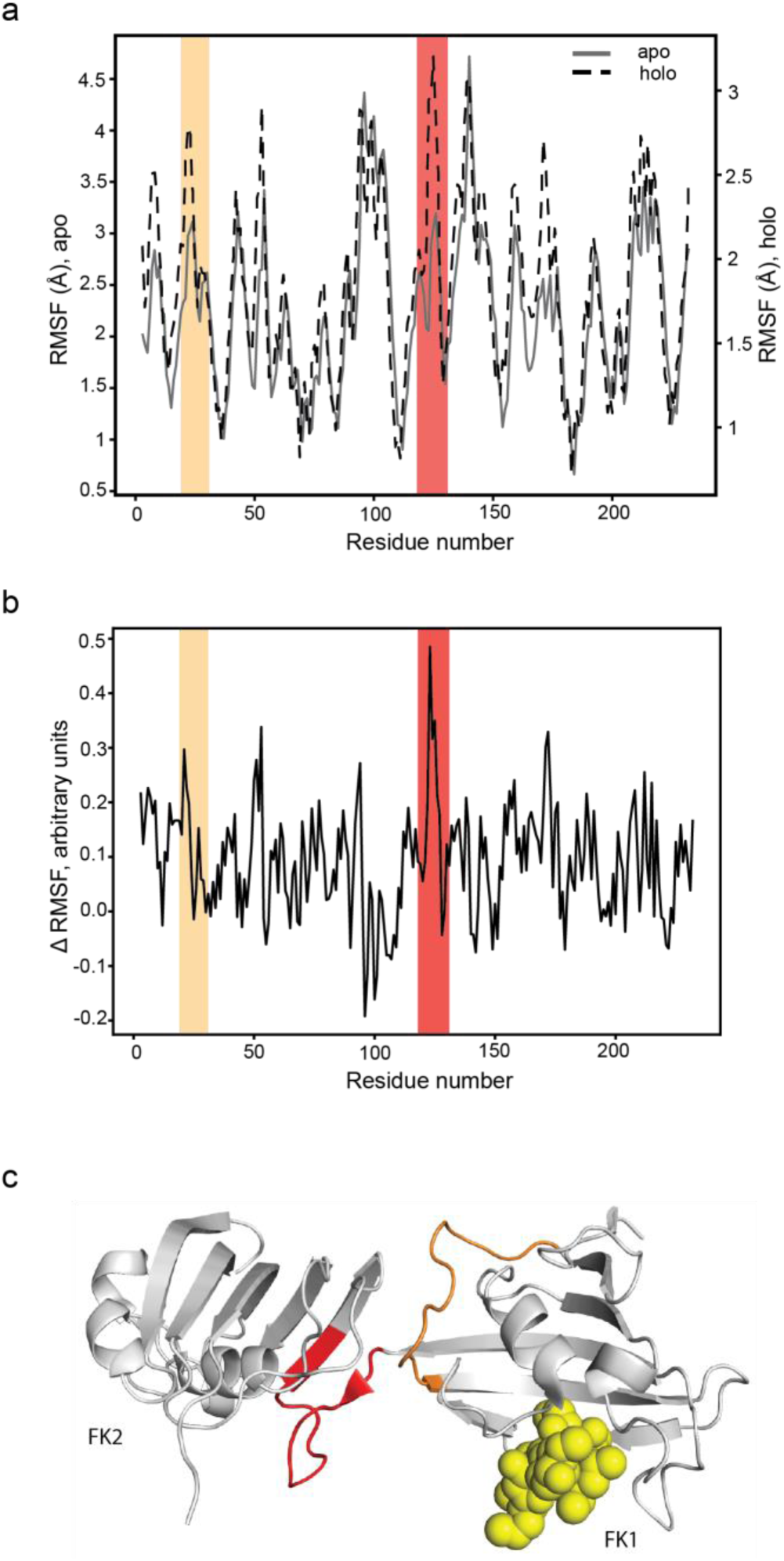
Comparison of AFDIP results with molecular dynamics simulations. (a) RMSF plots for apo and holo FKBP4. Parts of the sequence showing a shift in AFDIP are colored in red and orange. (b) Difference between the normalized RMSF values for holo and apo conformations. Parts of the sequence showing a shift in AFDIP are colored in red and orange. (c) FKBP4 structure (PDB: 1Q1C) with shifting peptides in AFDIP colored in red (flexible linker) and orange, and FK506 molecule colored in yellow.

### AFDIP for protein-metabolite interactions elucidation

Small molecule metabolites play essential roles in cellular physiology by interacting with protein targets, forming a complex protein-metabolite interaction (PMI) network that is crucial to decipher but challenging to study using traditional methods due to its scale and complexity^50^. Several chemical proteomics methods have been employed to study PMI, for instance TPP^50^ and LiP^51^. To further explore the potential of chemical proteomics in elucidating PMI networks, we decided to test AFDIP using acetyl-coenzyme A (AcCoA) as a model metabolite.

AcCoA is a central metabolite that plays a crucial role in numerous cellular processes^52^, including energy metabolism, lipid biosynthesis, and protein acetylation. As a key intermediate in the citric acid cycle and a major acetyl group donor, AcCoA serves as a critical link between carbohydrate, lipid, and protein metabolism, making it an ideal candidate for studying protein-metabolite interactions.

Altogether, 61,882 peptides belonging to 6739 proteins were quantified with time-AFDIP. Among the peptides with significant p<0.05 and ΔCoG>±0.5 h, 66 molecules were associated to proteins involved in CoA metabolism and production (Supplementary figure 1a). To prioritize the most relevant interactions, we ranked peptides based on the sum of their absolute ΔCoG values and log10(p) values. Notably, 8 out of the top 10 ranked peptides belonged to proteins with known roles in CoA metabolism and production, including ACACA, HADH, NAA50, NAA15, NAA10, HMGCS1, ACOT7, and NAA40. 7 out of top 10 potential targets found by Fisher’s combined probability test belonged to proteins involved in CoA metabolism and production.

Of particular interest was N-alpha-acetyltransferase 50 (NAA50), which had two peptides among the top 10 of our putative target list. Given that an X-ray structure of NAA50 in complex with AcCoA was available^53^, we found a 9.3 Å difference between the binding predicted by AFDIP and determined by crystallography (Supplementary figure 1b), while the maximum longitudinal distance of AcCoA ligand in crystal structure is 19.7 Å.

### Staurosporine: comparison with LiP and TPP

As the target identification data for HeLa cells treated with staurosporine are available from several chemical proteomics techniques, such as LiP^18^ and TPP^3^, we compared with them the time-AFDIP results (Figure 7a).

**Figure 7.**
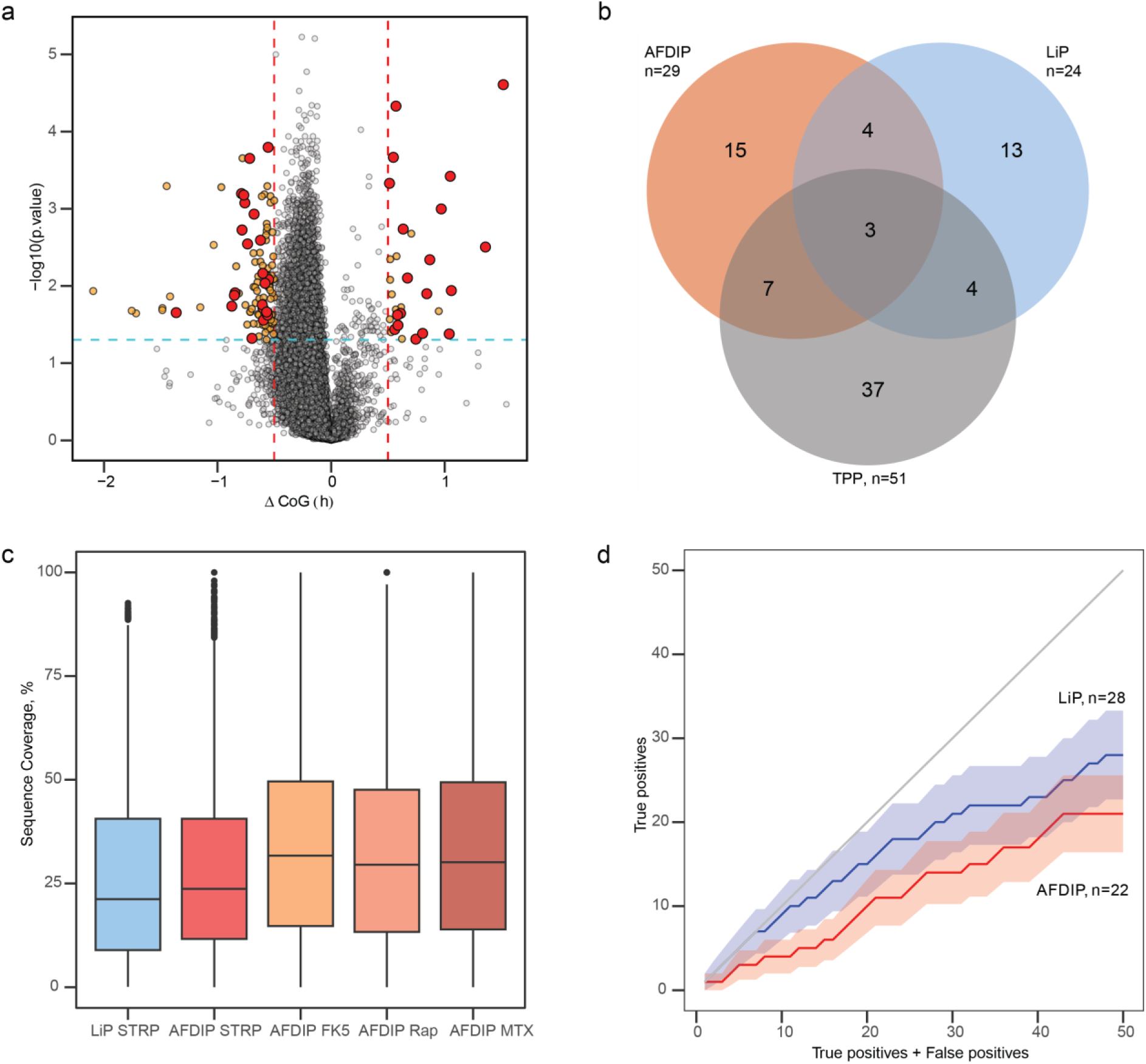
**Comparison of AFDIP, LiP and TPP in staurosporine analysis**. (a) Volcano plot for AFDIP showing all peptides having a significant center of gravity shift in orange (cut-off values are the same as in Figures 2-5) and peptides belonging to human kinases in red. (b) Venn diagram showing the kinase targets found with LiP^18^, TPP^3^, and AFDIP on the protein level. (c) Proteome sequence coverage for LiP ^18^ with staurosporine (light blue), and AFDIP with staurosporine, FK506, rapamycin, and methotrexate in different shades of red. (d) True positive rate evaluation for LiP^18^ (blue) and AFDIP (red) for staurosporine kinase target identification on the peptide level. The error bars represent standard deviation.

In total, 80,764 peptides belonging to 7859 proteins were quantified in time-AFDIP, compared to 5388 proteins in LiP Quant and 7678 proteins in TPP. The average sequence coverage (Figure 7c) in LiP was 26.5 %, while in AFDIP – 27.8 %, with a higher number of proteins identified. The average sequence coverage in AFDIP of FK506 was even higher, 33.7 %. The significantly deeper analysis in AFDIP than in LiP and higher average sequence coverage can be explained by the use of a single protease and the absence of semi-tryptic peptides that increases the burden of proof.

Overall, AFDIP identified with statistical significance 29 target candidates (‘pro-targets’), LiP – 24 pro-targets, and TPP – 51 pro-targets (Figure 7b). The TPP results combined with the overlaps between LiP and AFDIP were taken as “background truth” which was used to estimate the false positive and false negative rate of each proteolysis-based technique. The overlap between the AFDIP results and LiP was 7 pro-targets (24% of AFDIP proteins and 29% of LiP proteins), while the average expected random overlap was 1.3 proteins. Of the 7 overlapping proteins, 3 were also found among the TPP pro-targets. Thus the “background truth” dataset was composed of 55 pro-targets.

The overlap between the LiP pro-targets and the “background truth” dataset was 11 proteins (46%), while expected random overlap was 2.6 proteins. The corresponding data for AFDIP were 14 proteins (48%) and 3.1 proteins were expected to randomly overlap. This result indicates that AFDIP is at least as reliable in pro-target identification as LiP.

On the other hand, AFDIP did not outperform LiP in true positive rate identification (Figure 7d). But given the Poisson statistics of the data, the standard error bands (±σ) for the two techniques overlap to a large extent, making the difference between the techniques statistically insignificant.

## Discussion

AFDIP has demonstrated efficiency in drug target identification across multiple compounds and a metabolite, including MTX, rapamycin, FK506, staurosporine and acetyl CoA. For instance, AFDIP successfully identified DHFR as a primary drug target for MTX, multiple FK506-binding proteins as targets of rapamycin and FK506, and protein kinases as targets of staurosporine. This highlights the method’s capability to discern potential therapeutic targets effectively. The ability of AFDIP to distinguish between the modes of action of the drugs with similar but only partially overlapping targets, such as rapamycin and FK506, is particularly noteworthy. Although both drugs interact with FKBPs, they exhibit distinct affinities and trigger unique downstream signalling cascades. Rapamycin inhibits mTOR through FKBPs, whereas FK506 primarily affects the calcineurin/calmodulin pathway^49^. Our statistical analysis concluded that FKBP3 was a top target for rapamycin but not for FK506, and identified mTOR as one of rapamycin targets, underscoring AFDIP’s potential to capture subtle differences in drug mechanisms.

For staurosporine, while AFDIP identified fewer potential drug targets compared to TPP, it outperformed LiP and provided additional insights into protein structure that TPP does not offer. Furthermore, AFDIP achieved deeper proteome coverage, and showed higher average sequence coverage, indicating its potential as a complementary approach to existing chemical proteomics techniques. In addition to target identification, AFDIP has shown promise in elucidating binding sites. For all studied protein-small molecule complexes, the difference between the putative binding site predicted by AFDIP and the one identified by crystallography was under 10 Å. While the resolution may not be as high as that obtained with Cryo-EM or HDX MS, there is still considerable potential to extract meaningful insights into protein structure. For instance, investigating all shifting FKBP4 peptides from time-AFDIP with FK506 with MD suggested that negative CoG shifts could be correlated with allosteric changes in protein structure upon drug target binding. This effect would need to be investigated further using orthogonal techniques and studying allosteric modulators with AFDIP.

It is noteworthy that the structural resolution of AFDIP grows with the size of the protein or protein complex, which can be explained by the larger pool of identified peptides among which only top peptides are selected for structural elucidation of the binding site. This feature – higher resolution for larger complexes – seems to be unique for AFDIP, as competing structure-determination approaches, including X-ray crystallography, cryo-EM and HDX MS, show an opposite trend. Indeed, if the whole mammalian cell can be viewed as a single protein-dominated complex with a characteristic size of 100,000 Å (10 µm), AFDIP’s identification of the binding site of a drug with a <10 Å precision, or 10^-4^ on the relative scale, would be superior to any modern technique.

Summarizing, we believe that AFDIP has the potential to find its rightful place among the arsenal of proteome-wide approaches in chemical proteomics for target identification and characterization of drug-target interactions.

---

**Supplementary figure 1.**
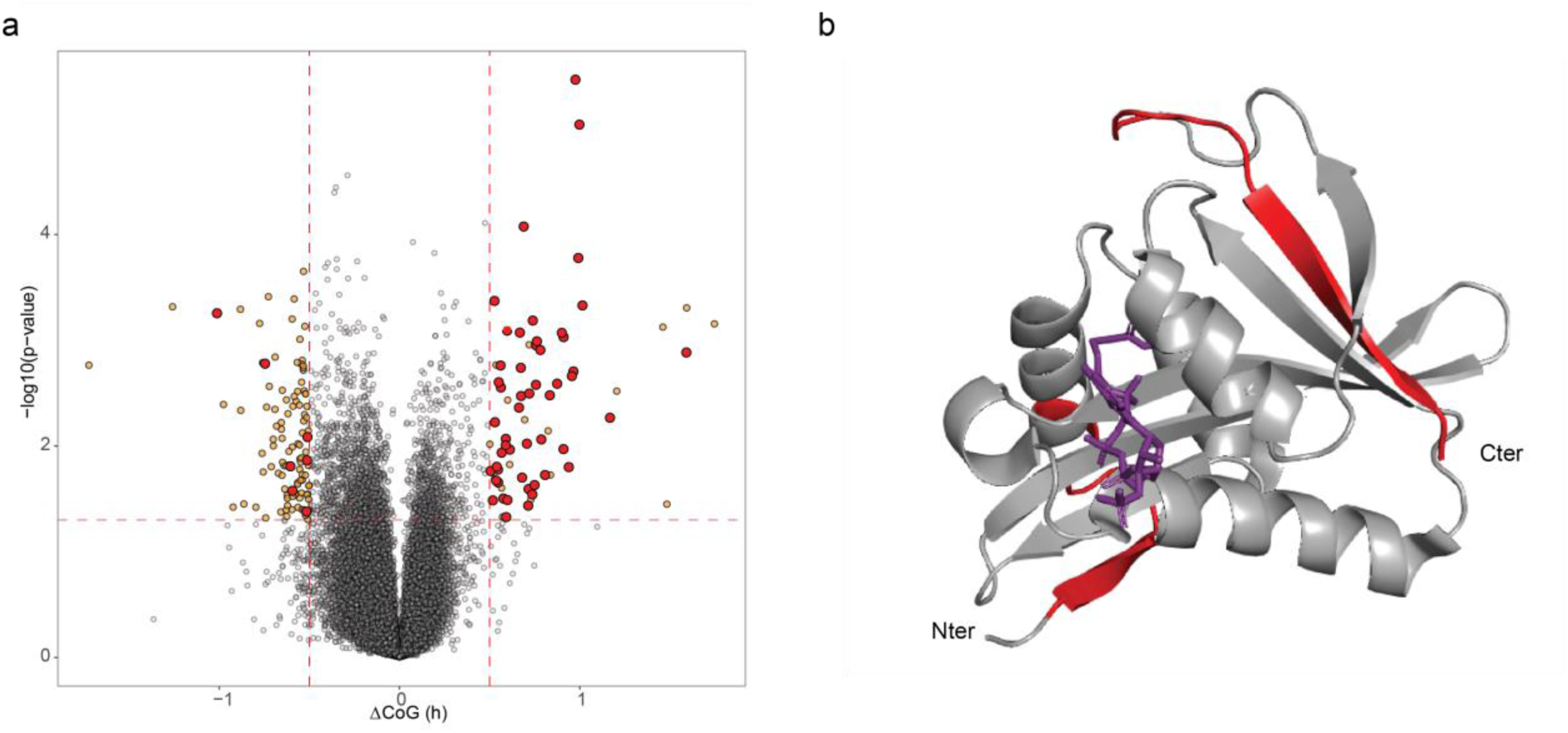
(a) Volcano plot for AFDIP showing all peptides having a significant center of gravity shift in orange (cut-off values are the same as in Figures 2-5) and peptides belonging to proteins involved in CoA metabolism and production in red. (b) NAA50 complex (PDB: 6WFN) with AcCoA. Shifting peptides used for CoM calculations colored red, AcCoA molecule in purple.

